# Interrogating mechanisms of ssDNA binding to a viral HUH-endonuclease by alanine scanning of an electrostatic patch

**DOI:** 10.1101/861070

**Authors:** Andrew Nelson, Kassidy Tompkins, Maria Paz Ramirez, Wendy R. Gordon

## Abstract

HUH endonucleases (dubbed “HUH-tags”) are small protein domains capable of forming covalent adducts with ssDNA in a sequence-specific manner. Because viral HUH-tags are relatively small, react quickly, and require no chemical modifications to their ssDNA substrate, they have great value as protein fusion tags in biotechnologies ranging from genetic engineering to single-molecule studies. One of the greatest assets of these tags is sequence-specificity to their unique, native *Ori* sequence *in vivo*, introducing the possibility of using multiple HUH-tags in multiplexed “one-pot” reactions. However, their mechanism of ssDNA sequence binding and specificity is poorly understood, and there is noted cross-reactivity between tags of closely related species. In order to understand the mechanism of ssDNA binding, we performed an alanine scan along a positively-charged patch of one such HUH-tag, replication-associated protein from Wheat Dwarf Virus (WDV Rep), and characterized the enzymatic activity in both the rate and extent of the reaction. In molecular beacon Stopped-Flow experiments, single point mutants of WDV showed a more than 60% decrease in reaction rate constant, and gel shift assays showed an almost complete lack of activity for some variants for single nucleotide ssDNA substitutions. In all, these findings help allow us to highlight key interactions in WDV-ssDNA binding, and we gain further insight into potential rational engineering of HUH-endonucleases to bind desired sequences of DNA.

## Introduction

The replication-associated domain of Wheat Dwarf Virus (WDV Rep) falls within a family of viral and bacterial phosphodiesterases known as HUH endonucleases (Yan and Fei 2007). In viruses, HUH endonucleases initiate rolling circle replication of the viral genome by recognizing the conserved nucleotide sequence of a ssDNA hairpin structure at its origin of replication and introducing a 5’-nick to the phosphate backbone. This nick is facilitated by the histidine-hydrophobic-histidine (HUH) motif of the protein, which coordinates a metal ion to stabilize a nucleophilic tyrosine. *In vivo*, this is a transient reaction, and the free 3’-end of the template strand re-ligates with the 5’-end at the end of genomic replication, releasing the Rep protein. In contrast, this reaction forms a robust phosphotyrosine adduct *in vitro*, and this protein-DNA conjugate is stable even in denaturing conditions (Chandler et al. 2013; Lovendahl, Hayward, and Gordon 2017; Heyraud-Nitschke et al. 1995). Thus, HUH endonucleases pose an attractive option for creating stable protein-DNA linkages in a variety of biotechnology applications, for example improving CRISPR-Cas9 precise gene editing (Aird et al. 2018) and tethering proteins of interest in DNA origami applications (Sagredo et al. 2016). Moreover, each HUH endonuclease recognizes a distinct sequence at its native origin of replication, resulting in a sequence-specific covalent adduct. Interestingly, HUH endonucleases derived from bacteria tend to be larger and less efficient at forming covalent adducts, but with more sequence specificity than their viral counterparts. Viral HUH endonucleases are small and form efficient covalent adducts, but their origin of replication sequences display more similarity, introducing the possibility of cross-reactivity. Understanding the molecular basis for single-stranded DNA recognition by viral HUH endonucleases offers the potential to engineer the viral HUH endonuclease to recognize desired DNA sequences, enhancing their multiplexing capacity. However, there are no crystal structures of HUH-endonucleases derived from viruses in complex with DNA. Thus we aimed to probe the requirements for ssDNA binding to the HUH-endonuclease domain from WDV by systematically mutating amino acids in a large, positively charged patch to alanine. The resulting variants were measured for effects on the kinetics of endonuclease activity, protein stability, and tolerance of nucleotide changes in the ssDNA substrate.

## Results and Discussion

We first identified electrostatic patches on WDV Rep using APBS electrostatics generation in Pymol and mapped them onto the recent crystal structure-pdb id: 6Q1M (Fig 1 A). We observed a large, positively charged patch in the vicinity of the catalytic residues (blue residues in Fig 1B comprising H59 and H61 involved in putative metal coordination and Y106, the site of phosphotyrosine bond formation) and reasoned that it might play some role in the binding and recognition of ssDNA. We mutated 11 amino acids (purple residues in Fig 1B) in this patch to alanine using site-directed mutagenesis, and these residues are colored yellow in the multiple sequence alignment of several viral HUH endonucleases shown in Fig 1C. It should be noted that several of the amino acids mutated are highly conserved while some are not. It is possible that these residues which are not conserved convey sequence specificity for WDV Rep which is exclusive from other geminiviral HUH endonucleases. Moreover, though the residues are clustered in a patch on the surface of the protein, they are spread across the protein in sequence space. Previous research has suggested that this patch is necessary for nuclear localization of other geminiviral Rep proteins, and it may serve bifunctionally to bind the negative charge on ssDNA as well <u>(Maio et al. 2019)</u>.

**Figure 1.**
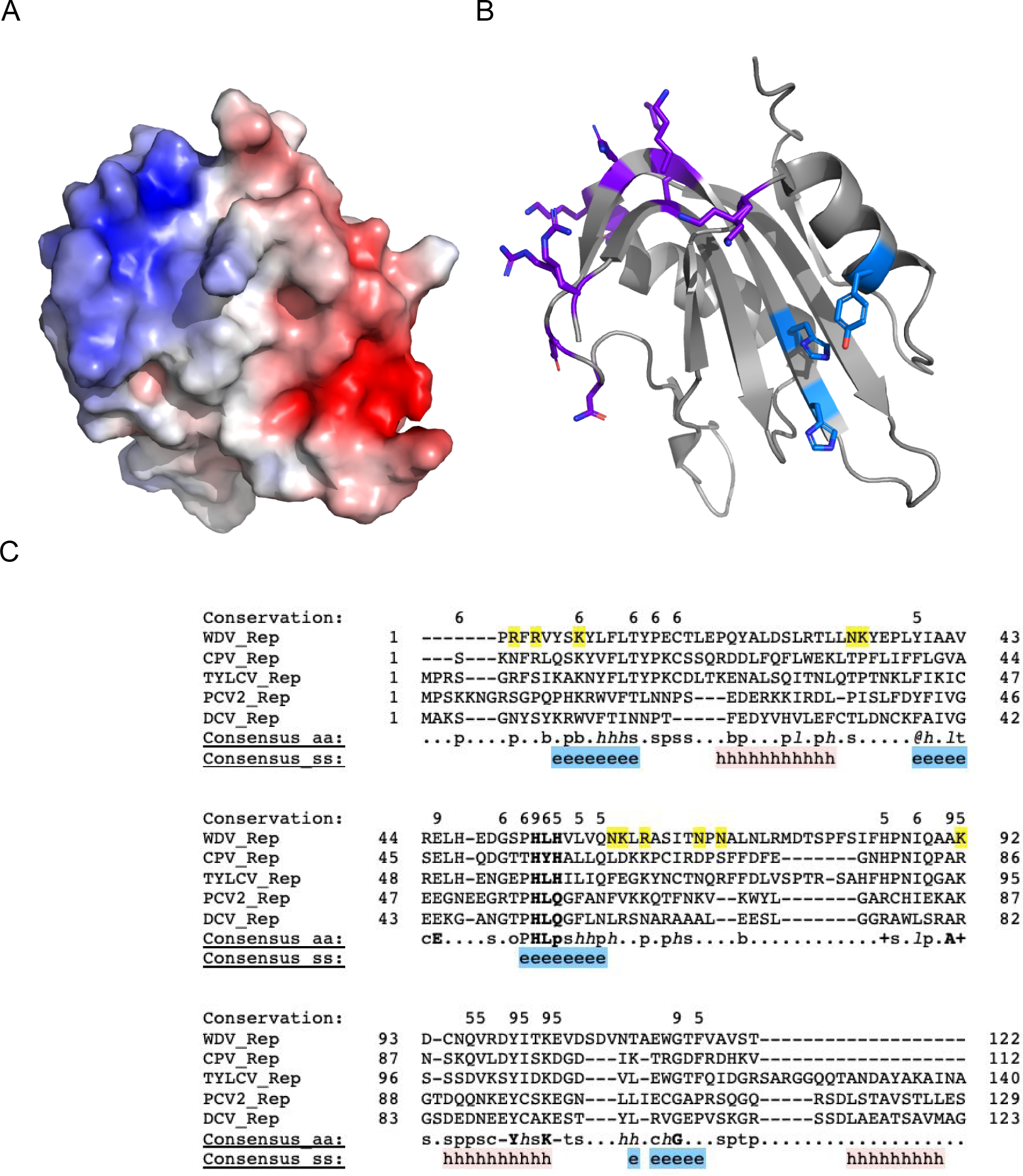
Alanine scanning of electrostatic patch. A. Surface representation of the WDV endonuclease with negatively charged regions colored red and positively charged regions colored blue. B. Side chains of amino acids of interest shown as ball and stick. Amino acids in the positively charged patch colored purple. Catalytic residues colored blue. C. Multiple sequence alignment of several viral HUH-endonucleases. Residues mutated in this study are highlighted yellow. Secondary structure depicted below alignment. Sequence conservation depicted by numbers above the sequence.

After expression and purification of these proteins using standard protocols, we used several assays to measure how the mutations affected aspects of HUH endonuclease activity. HUH endonuclease activity involves at least three steps: DNA binding, DNA cleavage, and DNA-adduct formation with the 5’ end of the cleaved DNA (Fig 2A). We first used a 26-nt DNA oligo containing the canonical origin of replication sequence in an SDS-PAGE gel shift assay to measure the extent of covalent adduct formed by the wildtype protein compared to the alanine mutants. The reactions were performed under “standard” HUH-tag reaction conditions (10x DNA, 50mM Hepes 8, 50mM NaCl, 1mM MnCl2; <u>Lovendahl et al. 2017</u>) at 37 °C for 30 minutes, and band volumes were quantified to determine covalent adduct formation. The data displayed in Figure 2B shows the efficiency of covalent adduct formation of all of the mutants with a 26-nt oligo harboring the “canonical” origin of replication sequence of geminiviruses. Surprisingly, the percent covalent adduct formation was not affected by mutation of the electrostatic patch. In order to assess if the kinetics of the WDV alanine mutant cleavage activity is altered as compared to the WT WDV, we used a molecular beacon, stopped-flow assay (Fig 2C). We devised a pseudo-first order scheme mixing 10x excess oligo substrate harboring the geminivirus canonical target sequence substrate flanked by a quencher and fluorescein mixed with WDV recombinant protein (standard buffer conditions). Single-turnover cleavage of the labeled oligo results in a positive readout of increased fluorescence monitored over the course of the reaction. We fit the kinetic curves using DataPro Viewer to a triple exponential (SI) (insert eq.) and displayed the fastest rate constant, k2, associated with cleavage. The other two rate constants may be partially describing binding and release of the substrate. All WDV alanine mutants showed decreased cleavage rates as compared to the WT WDV, which had a rate constant (k) of 0.28 s^−1^. Notably, the two most N-terminally positioned arginine mutants, R8A and R10A, had over a 50% decrease in cleavage. The three most C-terminally positioned mutants, N74, N76A, and K98A, along with the double mutant K67/98A, had very significant decreases in cleavage rates as well.

**Figure 2.**
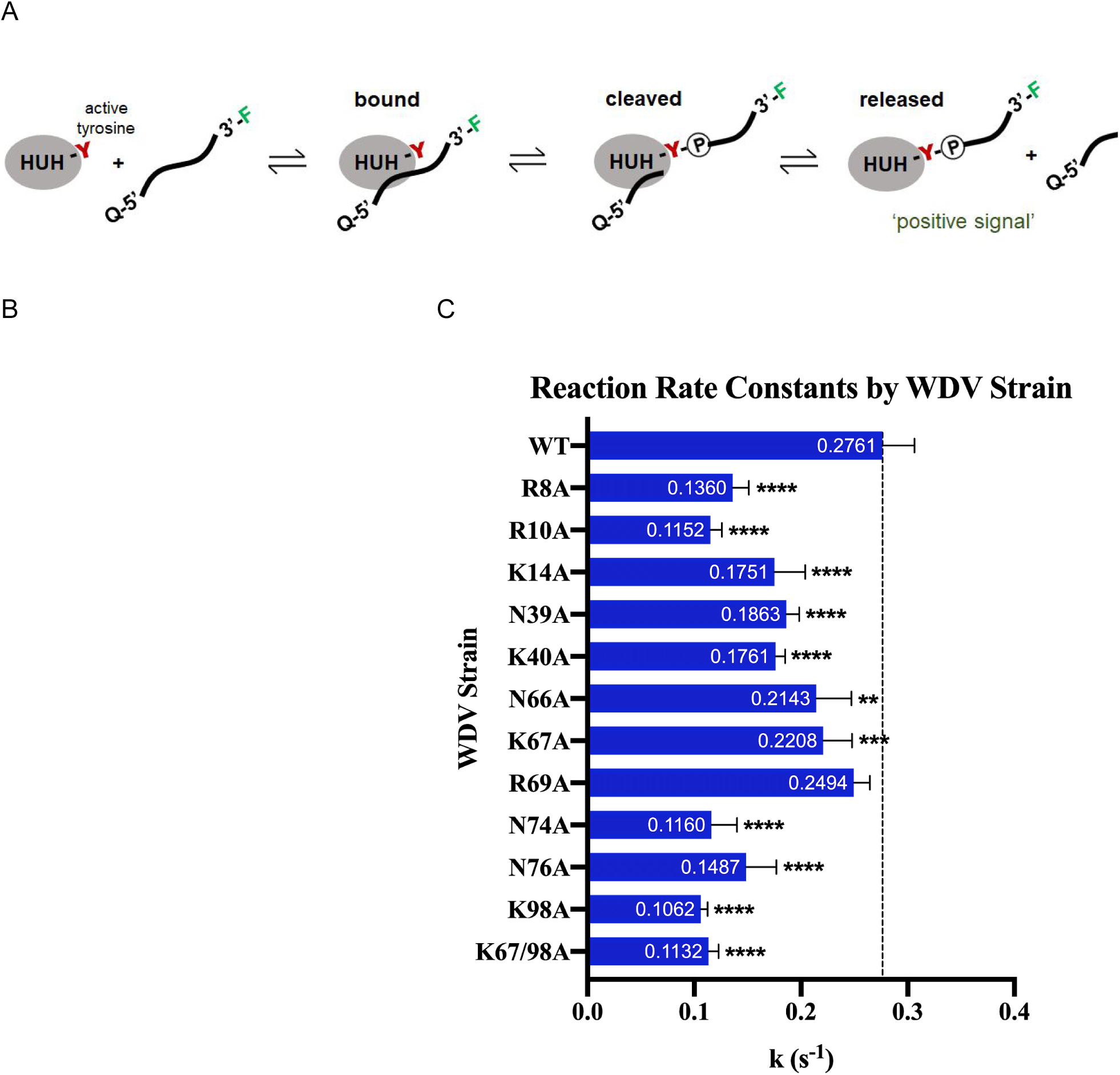
Mutation effect of WDV activity with canonical ssDNA sequence. A. Reaction scheme of HUH endonuclease formation of covalent adduct with ssDNA. B. Gel shift assay of WT and mutant WDV with ssDNA. C. Stopped-flow kinetics assay depicting DNA cleavage reaction of WDV and mutants with a molecular beacon ssDNA substrate. Statistical significance was calculated using ANOVA in Prism.

We next used Differential Scanning Fluorimetry to measure how the mutations affected the stability of the HUH-endonuclease to aid in deconvoluting the effects on DNA binding activity due to lost interactions versus destabilization of the protein. Shown is the average melting temperature of each mutant, computed from the second derivative curve of 4-8 replicate experiments (Fig 3). Several mutations showed up to 3 degrees decrease in melting temperature, and two mutants even showed enhanced stability. This suggests that the slower rates of cleavage of some of the mutants could, in part, be due to decreased protein stability.

**Figure 3.**
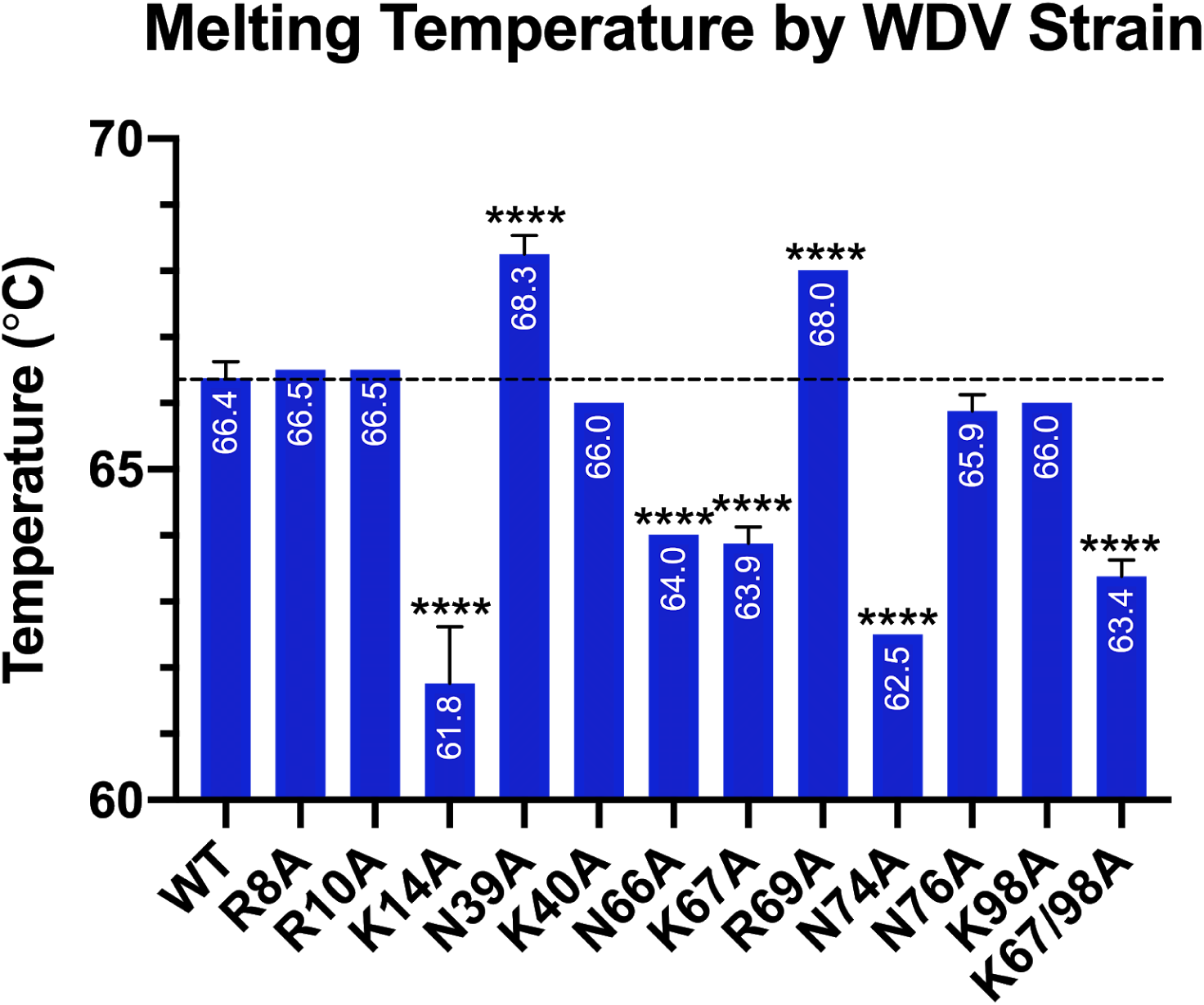
Stability of WDV and alanine mutants measured by Differential Scanning Fluorimetry.

**Figure 4.**
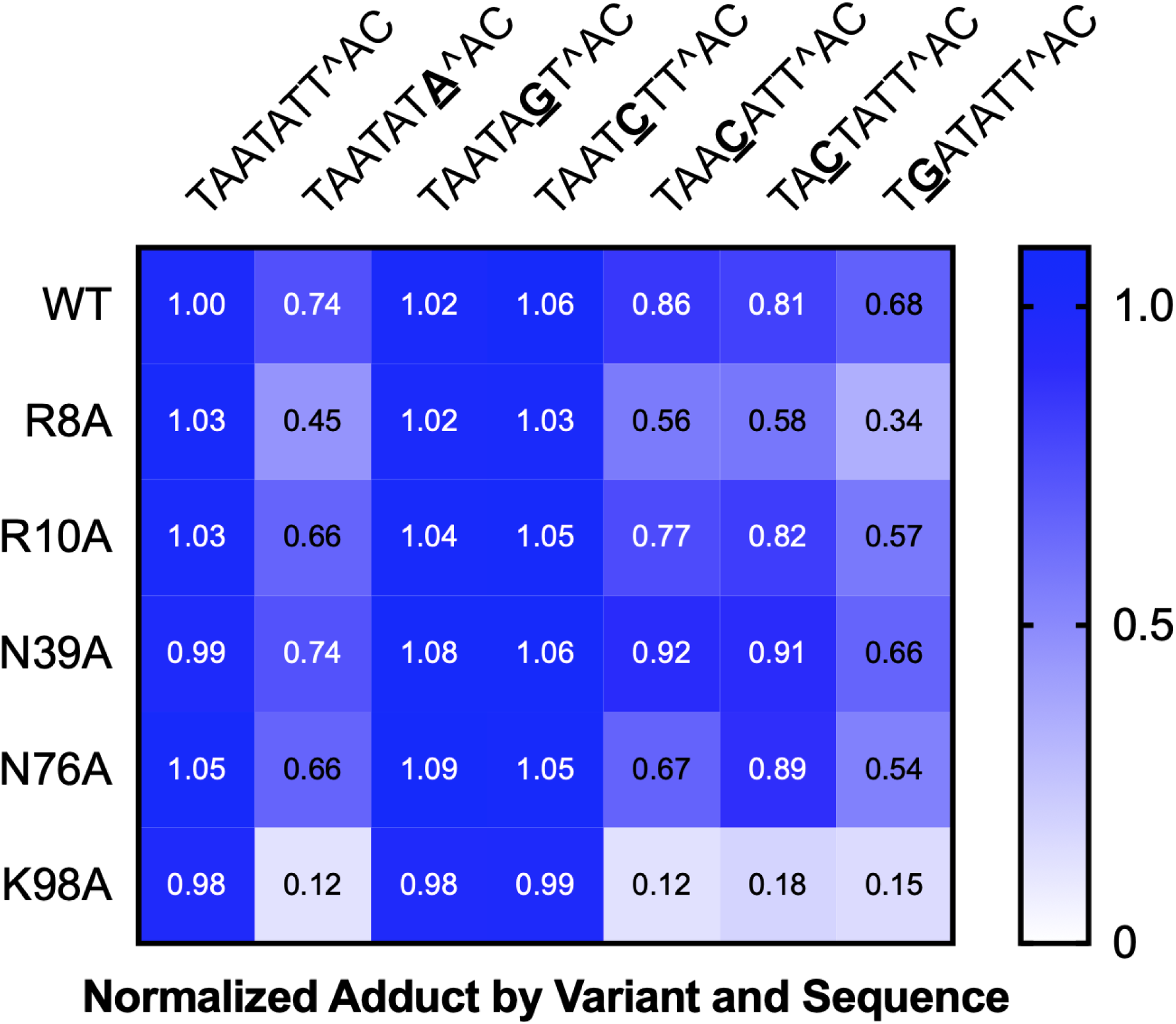
Covalent adduct formation by WDV and mutants upon reaction with altered ssDNA sequences. Gel shift assays were performed, and adduct formation was quantified using gel imaging software and normalized to the reaction of wildtype WDV with wtDNA sequence.

Finally, we measured the effect of mutations on binding non-canonical DNA oligo sequences to tease apart the potential roles of residues on DNA binding versus DNA specificity. Gel shift assays show the covalent adduct formation of each WDV variant for single nucleotide substitutions of the canonical 26-nt target sequence starting from the cleavage site and moving towards the 5’ (positions −1 through −6). Generally, most mutants had little to no change in covalent adduct formed in a five-minute reaction. However, the variants, R8A, R10A, N39A, and K98A showed noticeably decreased reactivity to nucleotide substitutions in positions −1, −4, −5, and −6.

Overall, the R8A and K98A mutants showed the most significant effects on activity with ssDNA with little to no changes to protein stability. Given that these are also highly conserved amino acids, it is likely that these residues are critical for binding DNA substrates, perhaps correctly positioning ssDNA for cleavage. Other, less conserved mutants showed differential effects on ssDNA binding activity, perhaps suggesting more subtle roles in dictating sequence specificity.

It should be noted that the substitutions which were chosen for the gel shift assay are not exhaustive. Future studies would benefit from current, next-generation sequencing methods to observe the full profile of sequence specificity for each variant. Furthermore, NMR studies may highlight WDV’s base interactions with DNA in a more sensitive, precise way than targeted mutagenesis. Inevitably, a crystal structure of WDV complexed to its ssDNA target would be most enlightening, and it would introduce the opportunity to rationally design the sequence-specificity of WDV. Our results provide a first step towards understanding the recognition of ssDNA by viral HUH endonucleases.

## Materials and Methods

### Homology Modeling and Mutant Design

We initially designed our mutations based on a homology model of WDV. WDV homology model was generated using SWISS-MODEL(Waterhouse et al. 2018) with the NMR structure of Rep of tomato yellow leaf curl virus (PDB: 1l5i(Campos-Olivas et al. 2002)) as a template. The generated structure with the lowest QMEAN score was downloaded and opened in PyMol (Benkert, Biasini, and Schwede 2011; DeLano 2002). During these studies, we obtained the crystal structure of apo WDV and used the structure in Figure 1. The surface electrostatics of the model was generated using the APBS electrostatics plugin in PyMol, and positively charged residues within the apparent electropositive patch were selected for mutagenesis.

### Primers

All mutagenizing primers were purchased from IDT. Sequences are as follows:

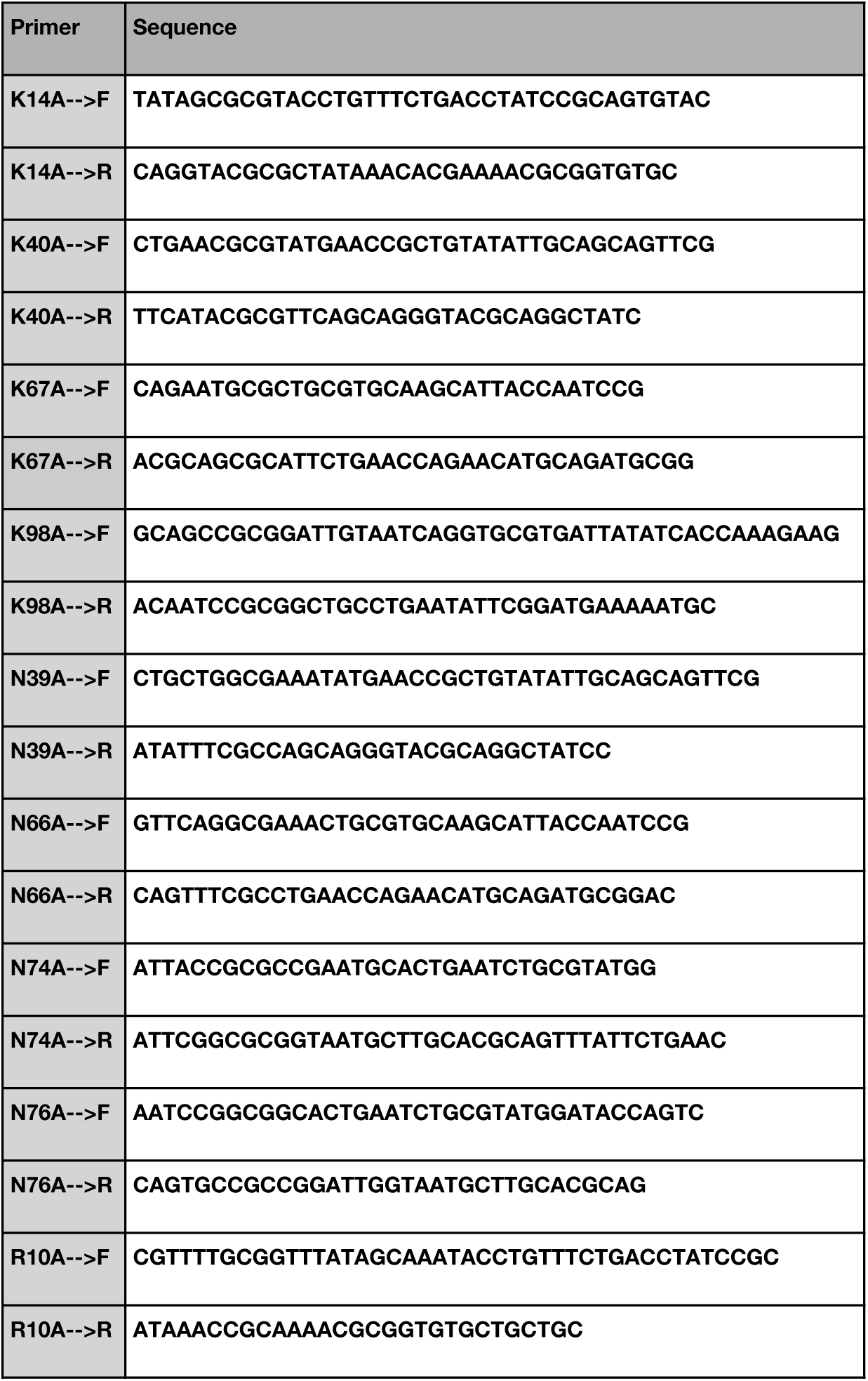
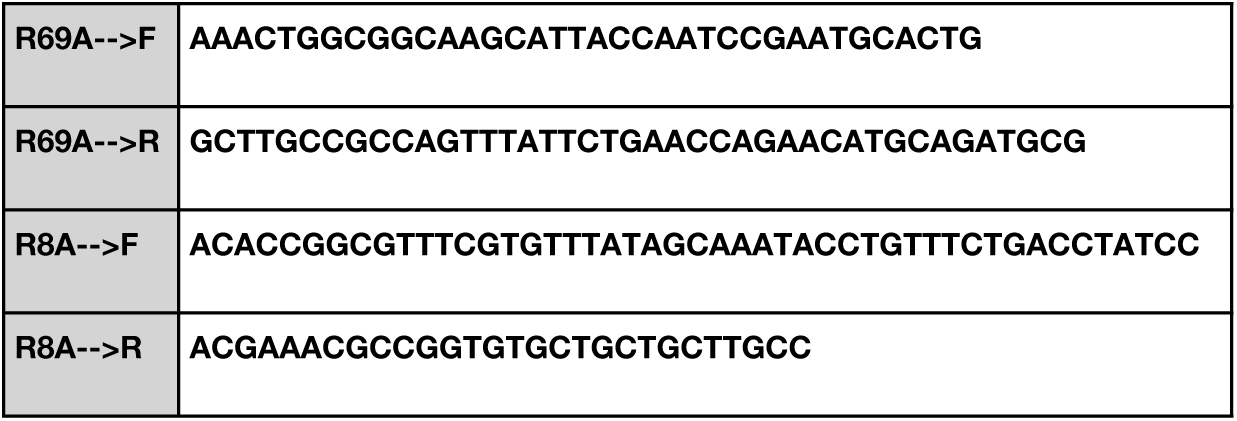

### Protein Expression

A pTD68 expression vector containing the coding sequence for residues 1-137 of WDV RepA (UniProt ID: P06847) attached to an N-terminal 6xHIS-SUMO tag was used for this study. Mutations to the RepA gene were introduced using In-Fusion primers (Takara) designed to contain the mutant Ala codon within the 15-bp homology arm (Supplemental For Primers). PCR amplification on these primers was done using CloneAmp HiFi PCR Premix (Clontech) and original template DNA was digested by DpnI. Sequence-confirmed, mutant plasmids were transformed into competent *Escherichia coli* (*E. coli*) BL21-DE3 cells and grown in 1L cultures of LB broth supplemented with 100 ng/mL ampicillin. Cultures were grown at 37°C to an OD_600_ of 0.8 before induction by 500 µM isopropyl β-D-1-thiogalactopyranoside (IPTG). Induced cultures were grown overnight at 18°C then pelleted by centrifugation at 4000 × G for 30 min. After resuspending in 6xHIS lysis buffer (50mM tris, 250mM NaCl, 1mM EDTA, pH 7.5), cells were lysed by sonication, and cellular debris was pelleted by centrifugation at 25000 × G for 30 min. Soluble protein was batch-bound to Ni^2+^-NTA resin and washed by 10 column-volumes of 6xHIS wash buffer (50mM tris, 250mM NaCl, 1mM EDTA, 30mM imidazole, pH 7.5). Protein was eluted in 6xHIS elution buffer (50mM tris, 150mM NaCl, 1mM EDTA, 300mM imidazole, pH 7.5), and imidazole was removed by size-exclusion chromatography through a Bio-Rad SEC70 column (Buffer: 50mM tris, 150mM NaCl, 1mM EDTA, pH 7.5).

### Differential Scanning Fluorimetry

SYPRO Orange dye (Thermo Fisher Scientific; 5000x in DMSO) was diluted 1:400 in experimental buffer (20mM HEPES, 500mM NaCl, pH 7.4). 30 µL of 2 µM proteins in experimental buffer were combined with 5 µL of the diluted dye in hard-shell, low-profile, 96-well, PCR plates (Bio-Rad). Plates were centrifuged at 4000 × G for 2 minutes before immediate analysis in a CFX96, real-time, qPCR machine (Bio-Rad). Proteins were held at 20 °C for 2 min, before the temperature was ramped to 90 °C at a rate of 0.5 °C/30sec. Plate RFUs were scanned every 30 sec using the FRET channel, and melting temperatures were determined from the second derivative of the RFU vs. Temperature curve.

### Circular Dichroism

Confirmation of DSF results were performed using thermal denaturation in circular dichroism (CD). Each purified protein was centrifuged at 14,000 × G for 10 minutes at 4 °C and the supernatant diluted to 0.35 mg/mL using PBS. Absorption at 222 nm was acquired with a Jasco J-815 spectropolarimeter, temperature controlled by a Peltier device. Changes in the absorption at 222 nm were acquired at a temperature gradient of 1 °C/min temperature intervals from 20–90 °C, and the characteristic ellipticity at alpha-helical wavelength (θ222) recorded. Molar ellipticity, [θ], was calculated using the following equation: [θ] = θ/(10*cl*) where *c* is the molar concentration of the sample (mole/L) and *l* is the path-length in cm. Molar ellipticity (with units of degrees, cm squared per decimole) was plotted against wavelength for the circular dichroism (CD) spectra. Ellipticity at 222 nm (θ222) was normalized, plotted against temperature, and fit by regression analysis in Sigma Plot (Systat Software, Inc.) using equations for two-state unfolding.

### Gel Shift Assay

In HUH reaction buffer (50 mM HEPES, 50 mM NaCl, 1 mM MnCl_2_, pH 8), 3 µM proteins were combined with 6 µM 26-nucleotide ssDNA sequences, and the reaction was allowed to proceed for 5 min at 37 °C. Reactions were stopped with 4x SDS loading buffer + 2% (v/v) β-mercaptoethanol and heat denatured at 95 °C for five minutes. Reactions were analyzed by gel electrophoresis on 4-20% polyacrylamide gels stained with coomassie blue for 30 min then de-stained in ddH_2_O overnight. Percent adduct was quantified using automatic band detection in Image Lab software (Bio-Rad).

### Stopped-Flow Kinetics

Molecular Beacon Stopped-flow HUH-tag Kinetics

Description: HUH-tag single turnover cleavage kinetics were characterized using a ssDNA molecular beacon assay on a standard stopped-flow instrument equipped with a fluorescence detector. The stopped-flow instrument (Applied Photophysics SX20) monochromater was set to 490 nm and 1.5 mm slit widths. Drive syringes were loaded with HPLC purified oligo substrate containing the geminivirus *ori* sequence and a 5’ iowa black-FQ quencher and a 3’ 3-6 fluorescein (`5’IABkFQ/CGTATAATATTACCGGATGGCCGCGC/36-FAM/`, IDT) or 100 nM HUH-tag each in reaction buffer, pH 8. A PMT detector (Hammamatsu) and 500 nm longpass filter (Edmund Optics) monitored cleavage of the substrate immediately following rapid mixing in the flow chamber for a duration of 2 minutes, and 1000 data points were collected in the logarithmic setting.

